# Indirect traumatic optic neuropathy after head trauma in adolescent male mice is associated with behavioral visual impairment, neurodegeneration, and elevated endoplasmic reticulum stress markers at acute and subacute times

**DOI:** 10.1101/2020.06.11.144766

**Authors:** Shelby M. Cansler, Fernanda Guilhaume-Correa, Dylan Day, Alicia Bedolla, Nathan K. Evanson

**Affiliations:** University of Cincinnati College of Medicine, Neuroscience Graduate Program; Virginia Polytechnic Institute and State University, Translational Biology, Medicine, and Health; Cincinnati Children’s Hospital Medical Center, Division of Pediatric Rehabilitation Medicine; University of Cincinnati, Department of Pediatrics

**Keywords:** traumatic optic neuropathy, head trauma, adolescent head trauma, ER stress, mice

## Abstract

Traumatic brain injury (TBI) results in a number of impairments, often including visual symptoms. In some cases, visual symptoms after head trauma are mediated by traumatic injury to the optic nerve, termed traumatic optic neuropathy (TON), which has few effective options for treatment. Using a murine closed-head weight-drop model of head trauma, we have previously reported in adult mice that there is relatively selective injury to the optic tract and thalamic/brainstem projections of the visual system. In the current study, we performed blunt head trauma on adolescent C57BL/6 mice, and investigated visual impairment in the primary visual system, now including the retina, using behavioral and histologic methods at multiple time points. After injury, mice displayed evidence of decreased optomotor responses illustrated by decreased optokinetic nystagmus. There did not appear to be a significant change in circadian locomotor behavior patterns, although there was an overall decrease in locomotor behavior in mice with head injury. There was evidence of axonal degeneration of optic nerve fibers with associated retinal ganglion cell death. There was also evidence of astrogliosis and microgliosis in major central targets of optic nerve projections. Further, there was elevated expression of endoplasmic reticulum (ER) stress markers in retinas of injured mice. Visual impairment, histologic markers of gliosis and neurodegeneration, and elevated ER stress marker expression persisted for at least 30 days after injury. The current results extend our previous findings in adult mice into adolescent mice, provide direct evidence of retinal ganglion cell injury after head trauma, and suggest that axonal degeneration is associated with elevated ER stress in this model of TON.

## Introduction

Traumatic brain injuries (TBIs) are diverse in cause, severity, location, and duration of symptoms. Although a number of models of TBI have been used in research settings, many facets of injury require further exploration. One of these avenues involves injury to the optic/visual system. Visual deficits are common after TBI, and can be due to damage to nearly any part of the brain, including the eye or optic nerve.[1] Severe and moderate deficits seen after TBI can include complete loss of vision, loss of visual fields, loss of color vision, or a decrease in visual acuity. Further, many symptoms may not be identified immediately, which can lead to missed diagnosis, especially in connection to the head injury.[2] Visual sequelae of TBI can also cause significant issues with recovery,[3] and can impact functional outcomes in children after even mild injuries.[4]

Among the pathophysiologic mechanisms of TBI-associated visual deficits is traumatic optic neuropathy (TON). TON is particularly associated with facial or frontal head impact, and can lead to severe visual impairment.[5] TON is often further classified as either direct (involving direct, penetrating injury) or indirect (injury due to transmission of force through the skull into the optic nerve, or due to secondary mechanisms like increased intraocular/intracranial pressure within the optic canal, axonal shearing, or neuroinflammatory responses). Indirect traumatic optic nerve damage, in particular, is a known comorbidity of traumatic head injury, in both adolescents and adults, with a reported incidence of between 2% to 5.2% of patients with closed head injury[5, 6]; however, it is likely that reported incidence rates are underestimated.[7]

While a direct optic nerve injury can be diagnosed by detecting optic nerve (ON) avulsion, ON transection, ON sheath hemorrhage (e.g. by imaging studies), indirect TON is generally more difficult to distinguish[8]. Fractures of the optic canal can be seen on CT if present, but often injury can only be detected by vision testing. [8] Optical coherence tomography (OCT) can also be a useful tool as it has revealed that indirect TON (iTON) results in significant thinning and deterioration of retinal layers at the time of injury and up to 35 years later, with retinal ganglion cell (RGC) death seemingly initiating subsequent deterioration of the other retinal layers.[2] Imaging of iTON within the brains of living human subjects, on the other hand, has only been possible through diffusion tensor imaging (DTI), thus limiting our knowledge of iTON. DTI has suggested that mild TBI can produce axonal injury in optic radiations [8-10] and the anterior thalamic radiations from the lateral geniculate body [9]. However, even advanced imaging modalities are limited in the information that they can provide.

Optic nerve injury has also been reported in rodent studies of TBI, including a couple done in adolescent-aged mice.[11, 12] Most studies of both human and animal optic nerve injury have focused on models relevant to direct TON (e.g. optic nerve crush or transection models), leaving both diagnosis and study of indirect TON less understood [13-15]. While these models provide valuable information into direct visual neuropathies, they are less relevant to indirect injury, which may be more common in patients with TBI [8]. Nevertheless, in recent years, several TBI models have reported optic tract injury along with other diffuse axonal injury [16-18], and, although many are repetitive TBI models [19-22], there are now at least three different proposed models that reproduce single-impact iTON [23-25].

Both animal and human studies of TON are typically conducted in adult populations (e.g., due to higher incidence in military personnel). As we and others have previously shown that outcome differences between mice separated by as little as two weeks of age arise after mild TBI in memory performance, mortality, and severity of brain pathophysiology [26-32] it is, therefore, likely that these differences are also present across age groups in the visual system. Moreover, studies on optic nerve injury have only recently begun to analyze central projections of the optic nerve like the LGN [17, 21, 33, 34], the SC [21, 23], and the supra-oculomotor nucleus and caudate [21] but none have been conducted in an adolescent population.

There are no evidence-based treatment options currently available for improving outcomes after TON.[35] A better understanding of the pathophysiology of TON is likely needed to develop more rational treatment approaches. For example, reactive oxygen species could be a potential target for treatment, based on research showing this as a potential pathologic mechanism.[25] Research in other optic neuropathy models further suggests that the unfolded protein response (UPR) may be involved in optic nerve degeneration.[36] Accordingly, we have developed a reproducible murine model of indirect TON associated with head trauma. We have previously reported degeneration in the optic tract, lateral geniculate nucleus, and superior colliculus within one week after TBI in adult male mice, but TON has not yet been studied in an adolescent population.[24] [26-32]In the current studies we expand our previous findings in adolescent mice, and include measures of retinal ganglion cell loss, changes to visual acuity, and changes in subcortical targets of the optic nerve both one week and one month after injury. We have added additional time points after injury to extend our understanding of the extent and duration of indirect TON. We further explore the hypothesis that endoplasmic reticulum stress (i.e., the UPR) may be a mechanism of early and lasting damage.

## Methods

### Animals

These experiments were performed in 6-week old adolescent [37] male C57BL/6J mice (Jackson Laboratories, Bar Harbor, ME). Animals were housed under a 14h:10h light:dark schedule in pressurized individually ventilated cage racks, with 4 mice per cage, and were given ad libitum access to water and standard rodent chow. All experimental procedures were approved by the University of Cincinnati Institutional Animal Care and Use Committee. Animals were allowed to habituate to the vivarium for one week prior to undergoing moderate head trauma and subsequent procedures.

### Traumatic Brain Injury

Closed-head TBI injury was performed by weight drop, as previously described.[38] In brief, mice were anesthetized using isoflurane (2-3%) and positioned under a metal rod (1.2 cm diameter; 400g), raised to 1.5 cm above the scalp in prone position on a .5 cm thick piece of cork board. Injury was produced by dropping the rod onto the calvarium with scalp intact, approximately above bregma (Fig. 1b). Compared to other weight drop methods, this approach uses a greater weight from a lower height, and appears to lead to injury to the optic nerve within the optic canal, but generally sparing other areas of the brain [24]. After injury, mice were immediately removed from the apparatus and allowed to recover. During recovery, obvious clinical seizures were observed and noted. Sham animals were subjected to anesthesia, then allowed to recover without undergoing the TBI procedure. Cohorts of mice were used for retinal (n= 9-12) and brain histology (n= 7-10) and protein analysis (n= 8-17), and for behavioral analysis (n = 8-12 per group). Figure 1 illustrates the timeline of experimental procedures performed and depicts location of head injury.

**Figure 1.**
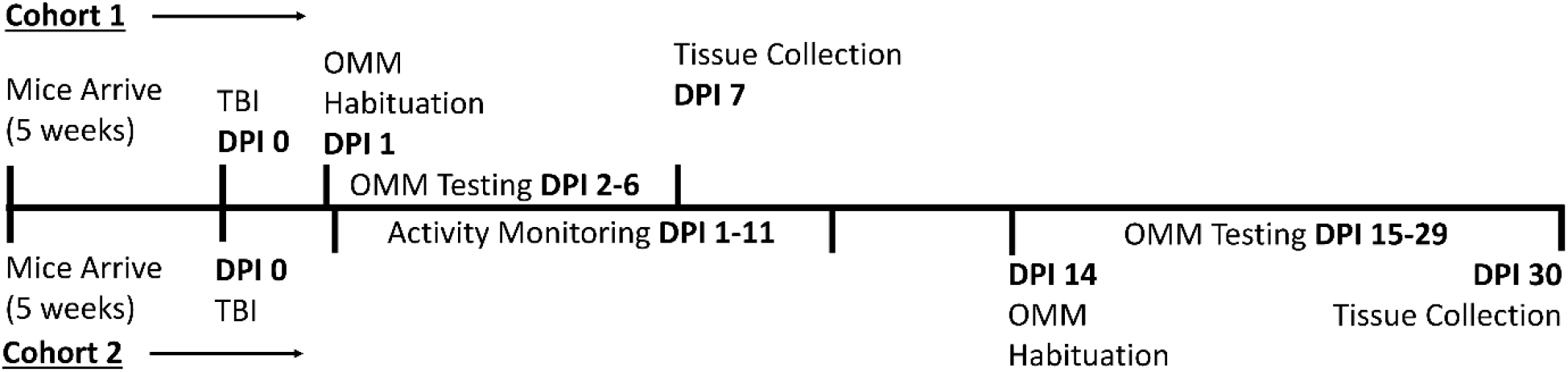
Experimental Timeline. Two cohorts of mice were utilized in this study. After arrival in the vivarium at 5 weeks of age, mice were allowed to habituate to the animal facility for one week prior to beginning experimental procedures. TBI/iTON by weight drop was performed on Day 0. Cohort 1 underwent only optomotor behavior testing, on days 2 – 6 post injury (DPI), followed by collection of eyes and brains at 7 DPI. Cohort 2, on the other hand, underwent activity monitoring on 1 – 11 DPI, optomotor testing on 15 – 29 DPI, and tissue collection of eyes and brains at 30 DPI.

### Behavioral testing

#### Circadian Rhythm and locomotion

Twenty-four hours after injury, animals were weighed and separated into individual cages with *ad libitum* food and water, but without any enrichment (e.g., no cotton bedding). They were then placed into activity monitors (San Diego Instruments PAS system, N=12; and Lafayette Activity Monitoring System, N=8). The systems were both set to record movements – beam breaks – for 24-48-hour time blocks. Every 24-48 hours, mice were weighed and the systems were reset. This was repeated for 7 days post injury. Beam breaks were binned into 1-hour periods for analysis. Activity monitors were used to analyze possible circadian rhythm shifts and overall activity levels.

#### Optomotor/Optokinetic Response (OKR) and Visual Acuity Testing

##### Optomotor Machine [OMM]

A machine was built to test optomotor reflexes, similar in design to the apparatus previously described[39] and illustrated in Figure 2A. The plexiglass case surrounding the mouse measures 20.2 inches high by 10 inch in diameter. The pedestal mice are placed on measures 7.4 inches long by 3 inches wide leaving roughly 3 inches of space between the mouse and the plexiglass on all sides. Sine-wave gratings are centered to extend 4 inches above and below the pedestal in order to encompass the mouse’s full frame of vision. The pedestal remains stationary as the plexiglass rotates on a circular stand. Speeds could be adjusted from 1 revolution per minute (rpm) up to 6 rpm and direction of spin could be switched between clockwise and counterclockwise.

**Figure 2.**
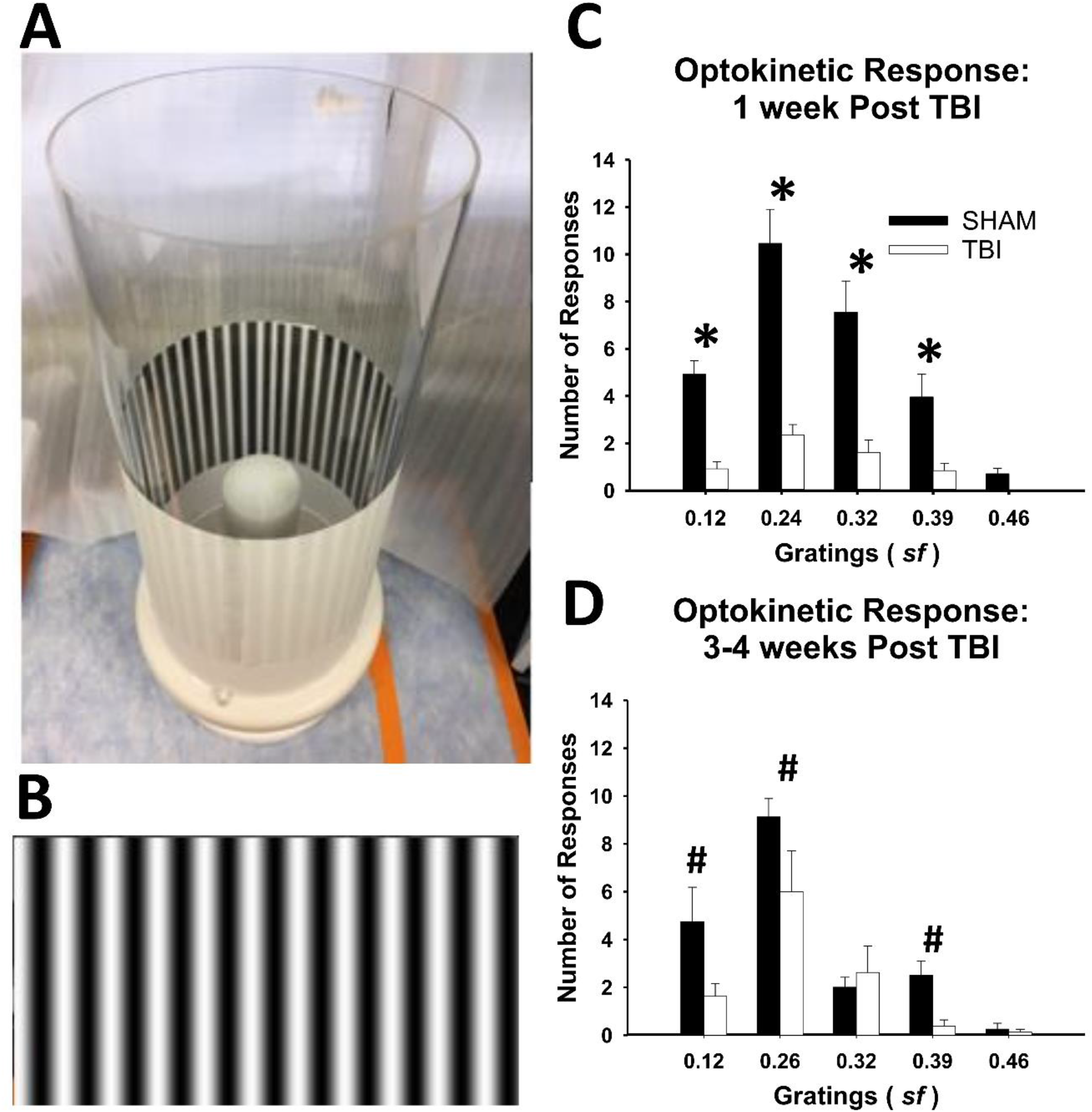
TBI/iTON mice have a significantly blunted optokinetic response 7 and 30 DPI. The number of responses were totaled for both left and right directions and are represented as mean ± SEM for each spatial frequency analyzed. A) Mice were placed on the white pedestal in the center of the machine and the plexiglass with the proper sine wave grating was placed around them. B) shows an example of a visual grating as used in this test. C) 7 DPI, iTON mice have significantly fewer responses at each grating and no responses at the highest grating of 0.46. D) 30 DPI there is still a deficit in responses, but this is no longer true at the 0.32 grating, which is within the reported optimal range of acuity for mice. * p <0.001, # p <0.005 vs sham at the same grating size.

##### Sine-wave Gratings

Sine-wave visual gratings were generated using the python package Psychopy[40] at varying spatial frequencies – 0.12, 0.26, 0.32, 0.39, and 0.42 cycles per degree (cpd) – as vertical, alternating, black and white lines. Each grating could be interchanged and attached to the inside of the machine (Fig. 2b). A series of 15 gratings ranging from 0.03-0.64 cpd were piloted (data not shown) based on reported ranges for mouse visual acuity and the aforementioned five were chosen based on optimal performance (i.e., consistently present optokinetic response) in normal C57BL/6J mice ([39, 41].

##### Testing

Order of subject, grating spin direction (i.e., clockwise or counterclockwise), and grating frequency were counterbalanced (such that animals were not always tested in the same order and gratings were not used in the same order in each animal. This was done to mitigate possible practice or timing effects of testing on the results) across 5 days. Animals were placed on a pedestal in the OMM without a grating and were allowed to habituate for 10 minutes on day 1 (either 24 hours after injury for the 7 days-post-injury (DPI) cohort or 15 DPI for the 30 DPI cohort) – after 5 minutes the machine was turned on and rotated. On day two of testing, animals were placed on a raised platform with the grating already in place (this was done to avoid unnecessary disturbances to the mouse and to prevent reactions to events that could be seen through the plexiglass cylinder) and were allowed to acclimate to the device for 5 minutes (with no rotation so as not to activate the response until the appropriate time). The OMM was then turned on and rotated in the first direction for 2 minutes at 2 rpm. The machine was then stopped, and a rest period of 30 seconds elapsed before it was turned on again for 2 minutes in the reverse direction (protocol adapted from [42]). This procedure was repeated 5 days in a row at the same time each day by the same experimenter (between 0830-1430). One mouse was tested at a time with only one grating tested per day. The machine was cleaned using an antiseptic wipe between each animal. A video camera placed above the device recorded the last minute of acclimation and the 4.5 minutes of the testing time.

##### Scoring

OKRs were tallied for the same amount of recording time for each animal (4 minutes of rotation time) from video footage collected on a SONY HDR-CX440 Handycam. Each video was viewed and scored by two trained reviewers who were blinded to experimental conditions. Reviewers were trained for at least 2-3 weeks on optomotor response criteria before scoring videos, and there were no significant inter-rater differences. Briefly, an OKR was defined as a rotation of the nose consistent with the speed and direction of the drum over >5° angle changes. The total number of OKRs for each grating were recorded and the average between the two reviewers was analyzed.

### Retina Dissection

Seven DPI animals were euthanized and eyes removed. After fatal overdose of pentobarbital, the left eye was proptosed, enucleated, and placed in ice cold 1xPBS. The retina was immediately dissected on ice, and flash frozen in dry ice for protein extraction (as described below). The right eye was removed after perfusion of the animal with 4% PFA by proptosing the eye and using curved scissors to sever the optic nerve for immunohistochemistry (IHC). The enucleated whole right eye was post-fixed in ice cold 4% PFA for 1 hour then submerged in 30% sucrose overnight. Eyes remained in sucrose at 4°C until ready for retina removal. In the 30 DPI cohort, our protocol was adjusted and both eyes were removed before perfusion rather than after with the same subsequent procedure for dissection of the left retina for freezing and post-fixation of the right eye for IHC– this did not appear to alter the appearance of tissues as can be seen between the sham conditions for 7 DPI versus 30 DPI tissue in Figures 3 and 4.

**Figure 3.**
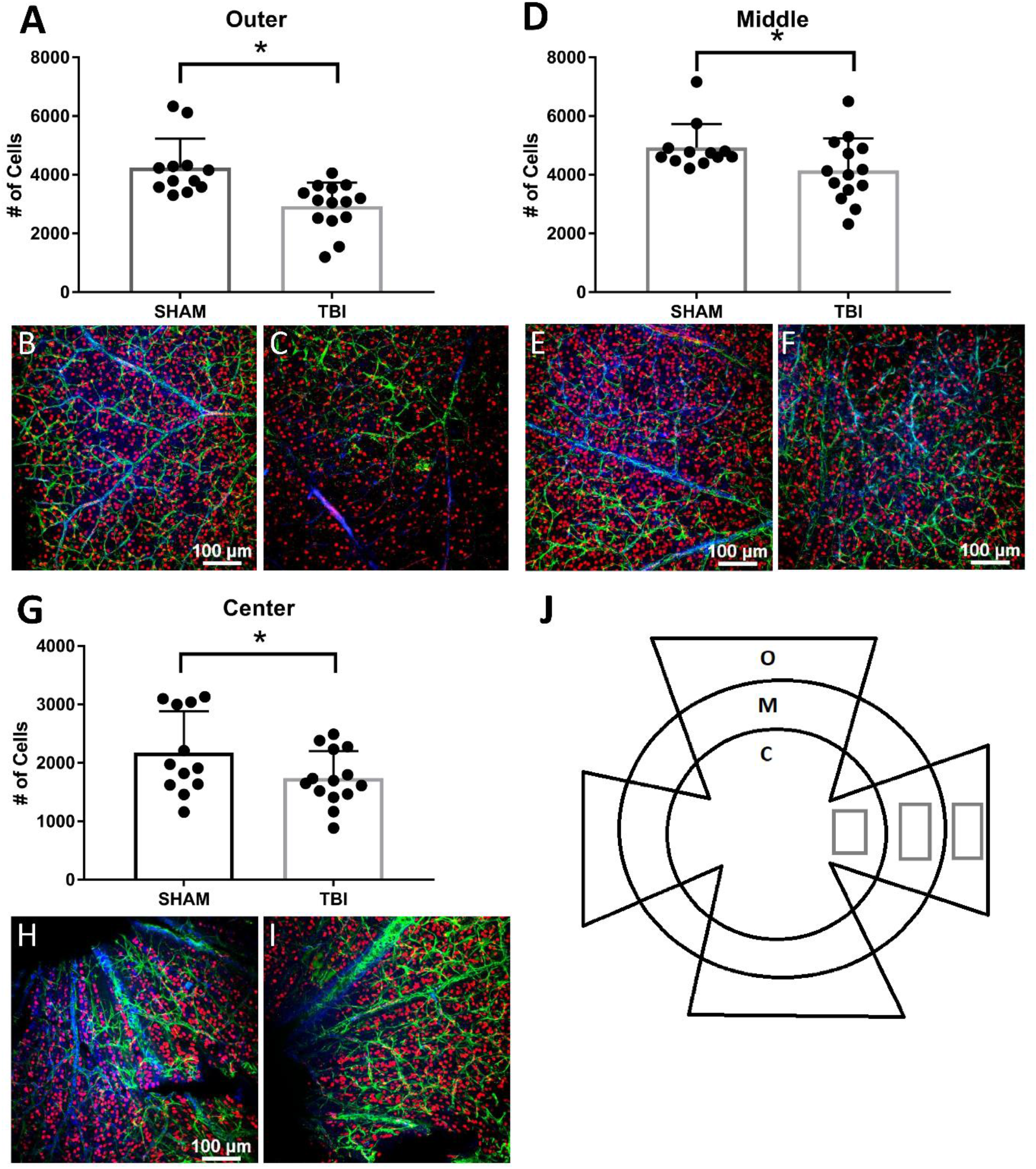
TBI/iTON mice have significantly fewer retinal ganglion cells throughout the retina at 7 DPI. Retinas of sham and TBI mice were immunolabeled with Brn3a (red), GFAP (green), and DAPI (blue). Brn3a labeled cells were counted in 3 zones of the retina (J). RGCs were decreased in all zones 7 DPI in iTON mice (A, D, G) Representative photomicrographs taken at 20x magnification of sham (B, E, H) and TBI/iTON mice (C, F, I). * p < 0.001

**Figure 4.**
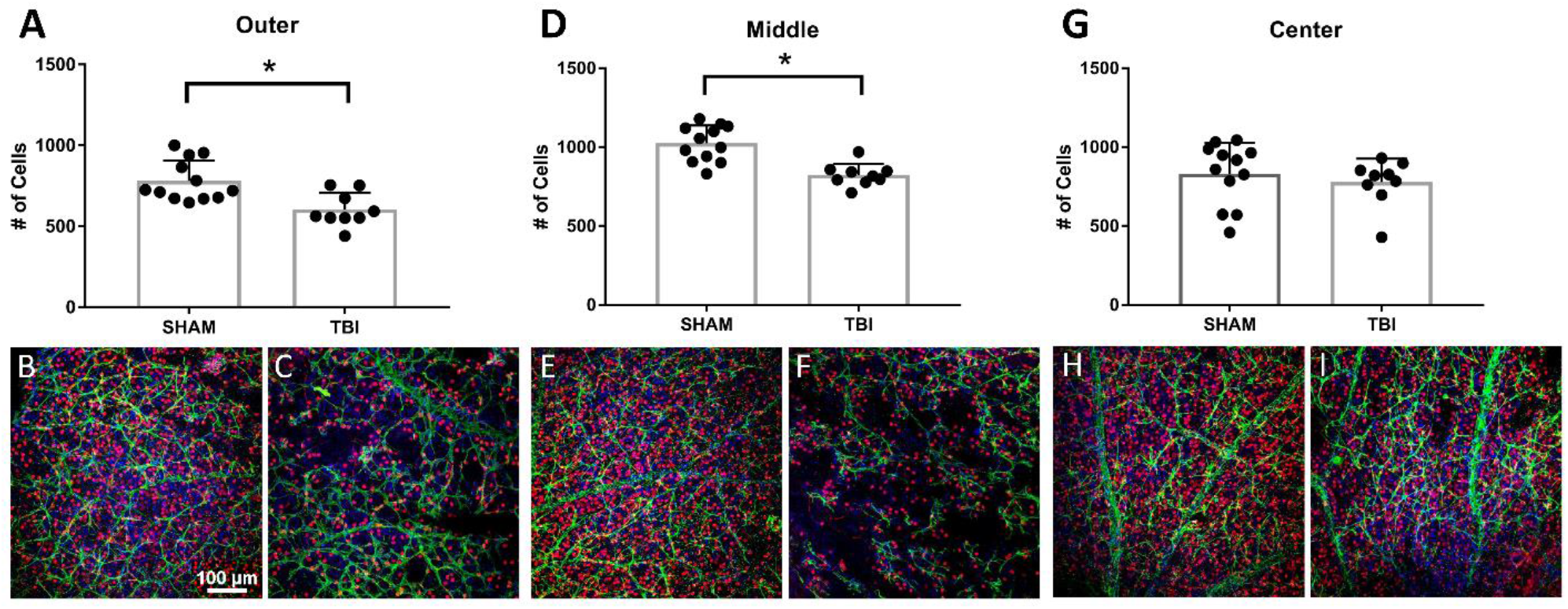
TBI/iTON mice have significantly fewer retinal ganglion cells throughout the retina at 30 DPI. Retinas of sham and TBI mice were labeled with Brn3a (red), GFAP (green), and DAPI (blue). RGCs were decreased in outer and middle zones of iTON mice (A, D) but not the center zone (G). Representative photomicrographs taken at 20x magnification of sham (B, E, H) and TBI/iTON mice (C, F, I). * p < 0.001

An adapted protocol from Ullmann, et al. [43] was used to dissect retinas. First, the eye was washed three times for five minutes in 1xPBS. In a petri dish with 1xPBS, the eye was punctured just above the ora serrata and microdissection scissors were used to cut along the cornea and iris until the corneal surface was removed. The lens, iris, and vitreous humor were then removed without touching the retinal surface of the interior eyecup. Finally, the sclera was gently pulled away from the retina and the retina was separated intact for whole-mount staining or protein extraction. Retinas (right eye only) used for IHC were stored at 4°C in 1xPBS in 1.7mL centrifuge tubes and stained within 48 hours of dissection. See below for detailed left eye protocol.

### Western Blots

After animals reached a deep surgical plane of anesthesia from Fatal Plus injection and before paraformaldehyde perfusion, the whole left eye was proptosed and curved scissors were used to sever the optic nerve in both 7 and 30 DPI mice. (As mentioned above, for the 7 DPI cohort, the right eye was removed after PFA perfusion for IHC; and in the 30 DPI cohort, the right eye was removed before perfusion and immediately submerged in cold 4% PFA for IHC.) In contrast to right eye fixation and washes before retinal dissection, left retinas were dissected without fixation in ice cold PBS on an ice bath immediately after removal. The rest of the dissection procedure remained the same. Unfixed retinas were placed into lysis buffer (20mM Tris-HCL pH 7.4, 2mM EDTA, 0.5mM EGTA, 1mM DTT, HALT protease/phosphatase inhibitor) and immediately put on dry ice. Retinas were then stored at −80°C until protein extraction. To extract retinal protein, retinas were homogenized with a pellet homogenizer for roughly 10 seconds. Samples were then sonicated in a cold water-bath for 5 minutes after which they were centrifuged at 3000 rpm for 20 minutes. Supernatant was removed and a BCA protein assay (Pierce BCA Protein Assay Kit; Thermo Fisher Scientific; cat # 23227) was used to calculate protein concentration.

Samples were prepared for western blot by measuring 20 μg of protein and adding β-mercaptoethanol and NuPAGE LDS 4x sample buffer (Invitrogen; cat # NP0007) and heating at 70°C for 10 minutes. Using SurePAGE 4-12% bis-tris gels (GenScript, Piscataway, NJ; cat # M00653), 40μl of protein sample was loaded into each well and run at 200V for roughly 1.5 hours in Tris-MOPS-SDS running buffer (GenScript; cat # M00138). Gels were then transferred via a wet transfer using Transfer Buffer Powder in 1xTBST (GenScript; cat # M00139) for 1 hour at 30V onto Amersham Hybond P 0.45 PVDF membranes (GE Life Sciences, Pittsburgh, PA; cat # GE 10600029). After transfer membranes were rinsed in TBST and blocked for one hour in either 5% milk (PERK, Caspase-3, ERO1L) or 5% BSA (βActin, IRE-1α, CHOP, and PDI).

Membranes were then incubated 24-48 hours at 4°C in primary antibodies – PERK (1:500 anti-Rbt; Cell Signaling Technologies [CST], Danvers, MA; cat # 3192S), IRE-1α (1:500 anti-Rbt; CST; cat # 3294S), PDI (1:2000 anti-Rbt; CST; cat # 3501S), CHOP (1:500 anti-ms; CST; cat # 2895S), Caspase-3 (1:5000 anti-Rbt; CST; cat # 9662S) ERO1-L (1:1000 anti-Rbt; Thermo Fisher Scientific; cat # 702709) and β-Actin (1:3000 anti-ms; CST; cat # 3700S). After incubation, membranes were rinsed in TBST then incubated in respective anti mouse HRP (CST; cat # 7076S) or anti Rabbit HRP (CST; cat # 7074S) for 1 hour at room temperature. Finally, membranes were washed and incubated in Pierce ECL PLUS Western Blotting Substrate (Fisher Scientific; cat # 32132) for 1-5 minutes after which they were placed between transparency film and taken to a dark room for (CL-XPosure Film; Thermo Scientific; cat # PI34090) exposure (times ranged from 2 minutes-4 hours). Films were scanned and converted to 8-bit TIFs for analysis using Image-J software.[44] Using the Gel Analysis tool peaks across protein bands were recorded. Each protein band was normalized to β-Actin.

### Histology

For histologic analysis of brain tissue, mice were euthanized using Fatal Plus® 7 days and 30 post-injury as previously described.[26] Sections were stained using Fluoro-Jade B or fluorescent immunohistochemistry. Fluoro-Jade B (Histo-Chem, Jackson, AR; cat# 1FJB), a marker for degenerating neurons and axons,[45] was used according to the manufacturer’s directions. After staining, slides were allowed to air dry completely in the dark and left un-coverslipped to avoid high background (Electron Microscopy Sciences, Hatfield, PA; cat# 13512). Slides were stored in a slide box and kept in the dark until imaging.

### Immunofluorescence

Primary antibodies used for immunofluorescence on brain sections were polyclonal rabbit anti-glial fibrillary acidic protein (GFAP; DAKO, Santa Clara, CA; cat # Z0334; RRID AB_10013382) and ionized calcium-binding adaptor molecule 1 (Iba-1; Synaptic systems, Goettingen, Germany; cat# 234003, RRID AB_10641962), both at 1:2000 dilution. Brain sections were washed in PBS, and then incubated in blocking solution (PBS with 0.1% bovine serum albumin, 0.4% Triton X-100) for 1 hour. Following this, sections were incubated overnight at 4°C with primary antibody in blocking solution. On the second day, sections were washed, then incubated with Cy-3 conjugated secondary antibody (Jackson Immunochemicals, West Grove, PA; cat# 711-165-152, RRID AB_2307443) at 1:500 dilution for 1h, covered, at room temperature. Sections were mounted in PBS with 1 ml of 5% gelatin added, allowed to dry in the dark, rinsed in water and allowed to dry again. Slides were coverslipped using antifading polyvinyl alcohol mounting medium (Sigma-Aldrich, St. Louis, MO).

Primary antibodies used for immunofluorescence on whole-mounted retinal sections were brain-specific homeobox/POU domain protein 3A (Brn3a; Millipore; cat #MAB1585; RRID:AB_94166) at 1:1000 and GFAP (1:1000). DAPI staining was achieved with Vectashield Antifade Mounting Medium with DAPI (Vector Laboratories; cat # H-1200; RRID: AB_2336790). All steps were performed in a centrifuge tube to avoid contact and handling of the retina. Retinas were washed in PBS, incubated in 0.3% H_2_O_2_ for 20 minutes, washed again, then permeabilized in 0.5% Triton X-100 for 15 minutes at - 80°C. Tissue was thawed, washed in fresh permeabilization solutions for 20 minutes, then blocked (2% TX-100, 2% BSA, and 5% normal goat serum in PBS) for 1 hour at room temperature. Retinas were then incubated in Brn3a primary antibody 48-72 hours at 4°C. After primary incubation, retinas were washed in 0.5% TX-100 and incubated for 1.5 hours in anti-mouse biotinylated secondary antibody (1:400; Vector Laboratories; cat # BA-9200; RRID: AB_2336171). Washes were performed followed by treatment in VECTASTAIN Elite ABC HRP Kit (1:800; Vector Laboratories; cat # PK-6100; RRID: AB_ 2336817) for 1 hour. Following ABC, retinas were incubated in Cy3 streptavidin (1:500; Invitrogen, Grand Island, NY; cat # 434315) for 2 hours at room temperature covered. Retinas were then washed, incubated in blocking solution for 30 minutes, and left in GFAP primary antibody 24-48 hours at 4°C. After incubation in the second primary antibody, retinas were washed, incubated in anti-rabbit Alexa 488 (1:500; Invitrogen; cat # A11034) for 2 hours at room temperature, washed, and then mounted.

Retinas were mounted onto positively charged slides pretreated with Gattenby’s Solution (0.5% gelatin, 0.05% chromium potassium sulfate dodecahydrate (CrK(SO_4_)_2_ · 12H_2_O; Fisher Scientific, Grand Island, NY; cat # C337-500) in ddH_2_O. To do this, whole retinas were transferred to slides in 1xPBS and 4-5 radial cuts were made to create a “petal” or “cross” shape allowing the retina to lie flat on the slide.

### Image analysis

Photomicrographs of all brain slides were taken by a blinded observer, using an Axio lmager Z1 microscope with an Apotome (Leica Microsystems, Buffalo Grove, IL). All slides were photographed using the same exposure time and magnification within planned comparison groups. FJ-B stained slides and retinal whole-mounts were viewed on a Nikon C2 Plus Confocal Microscope (Nikon Corporation, Melville, New York). FJ-B slides were imaged using the same fluorescence intensity (on the FITC channel) and background reduction within comparison groups and magnifications. Retinal slides were imaged using three channel wavelengths for TRITC-FITC-DAPI; because these images were only used for cell counting, fluorescence and background reduction were corrected for each image to allow the best representation of cells within each region of interest. Retinas were separated into three zones (peripheral, mid-peripheral, and center) with 3-5 images (depending on how many times a retina was cut) taken of each zone for each retina (Fig. 3j). Images within each zone were averaged to get a representative RGC count for that zone – this number was used in statistical analyses.

Image analysis and quantification were performed using ImageJ software[44] for GFAP and FJ-B images or NIS elements software (Nikon, Melville, NY) for Iba-1 and Brn3a. For GFAP and FJ-B images, mean fluorescence intensity was measured in multiple non-overlapping samples within the relevant region. For Iba-1-stained tissue, images were thresholded, and automated soma perimeter measurements were taken for all cells above threshold within the region of interest. Comparisons were performed only within a single region. Brn3a cell counts were acquired using NIS elements automated detection settings and were thresholded so that only complete, non-overlapping cells were counted.

### Statistical analysis

Statistical analysis was performed using the SigmaPlot software package (Systat, San Jose, CA) – 3-way comparisons were analyzed using Statistica (TIBCO, Palo Alto, CA). Weight gain (treatment x day), activity monitor (treatment x day) and OKR data (treatment x grating) were analyzed using 2-way analysis of variance (ANOVA) with repeated measures. Significant effects were further analyzed using the Holm-Sidak post-hoc method. Histologic and protein measures were analyzed using unpaired t-test. Data were transformed if needed so as to not violate normality and equal variance assumptions. Significance was set a priori at p < 0.05. In graphs, data are represented as mean +/-standard error of the mean.

## Results

### TON Injured mice have a blunted optokinetic response

Injured mice tested from two to six DPI displayed a significantly reduced number of optokinetic responses compared to their sham counterparts (main effect of injury: F_1, 123_ = 45.5, p < 0.001). There was also a predicted main effect of spatial frequency (F_4,123_ = 32.8, p < 0.001). Post-hoc analyses revealed significantly reduced responses for TON animals at all spatial frequencies (p < 0.001) suggesting that there is impairment in the OKR and visual acuity (Fig. 2c).

Thirty DPI animals performed similarly with main effects of injury (F_1,79_ = 9.9, p = 0.007) and spatial frequency (F_4,79_ = 22.6, p < 0.001). Post-hoc analyses revealed that only 0.12, 0.24, 0.39 spatial frequencies were significantly reduced in TON injured mice compared to sham (respectively p = 0.04, p = 0.04, and p = 0.004; Fig. 2d).

### There is significant retinal ganglion cell death in TON mice throughout the retina

Retinal ganglion cell counts were taken 7 (Fig. 3) and 30 DPI (Fig. 4) to determine whether this injury was localized to the optic nerve. To acquire an accurate sampling of RGC densities throughout the retina, counts were taken in three zones as explained in our methods section (Fig. 3j). At seven DPI there were significantly fewer RGCs in TON injured (n=14) mouse retinas compared to shams (n=12) in the peripheral (t(24) = 6.3, p < 0.001), mid-peripheral (t(24) = 8.2, p < 0.001), and central regions (t(24) = - 4.0, p < 0.001). Cell counts were also significantly reduced in TON (n=12) compared to sham (n=9) mice 30 DPI but only in the peripheral (t(19) = 3.5, p = 0.002) and mid-peripheral (t(19) = 4.7, p < 0.001) quadrants but not in the center (t(19) = 0.9, p = 0.4). TON mice also had significantly elevated Caspase3 seven DPI (t(21) = −5.8, p < 0.001) suggesting active cell death occurs up to at least 7 days after injury. This elevation was no longer significant at 30 DPI (t(22) = −0.3, p = 0.7).

### Activity but not circadian rhythm is affected in this model of TON

Injured mice were significantly less active than their sham counterparts (F_1,43_ = 4.7, p = 0.04; Fig. 5g). Despite major optic pathways between RGCs and brain regions controlling circadian activity, circadian pattern of activity (i.e., more active at night) fluctuated similarly in injured and sham mice resulting in a significant effect of light vs. dark activity (F_1,43_ = 147.5, p < 0.001). There was no effect of DPI (F_1,43_= 1.0, p = 0.46) and reduced activity did not improve across the testing days.

**Figure 5.**
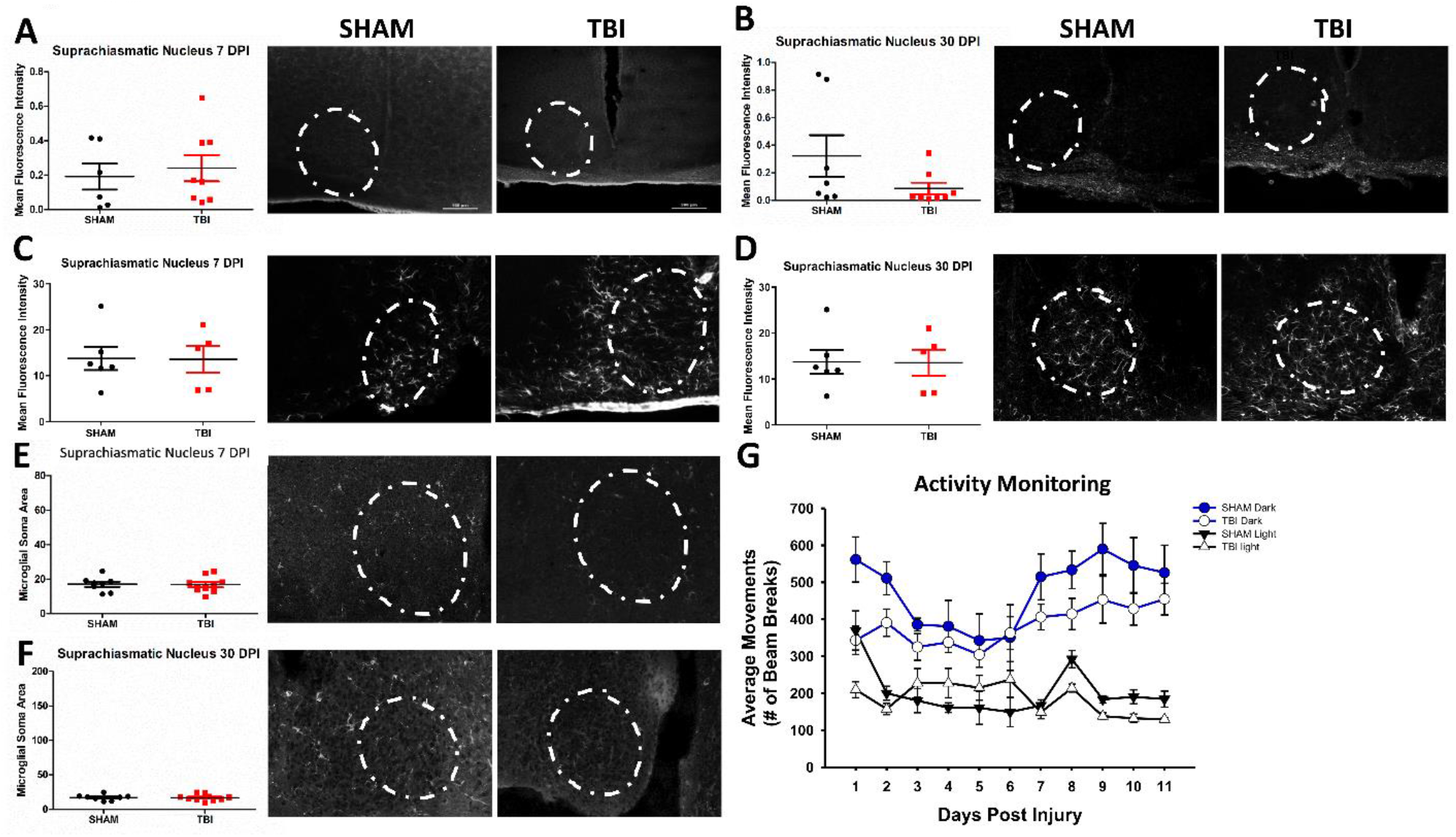
There are no circadian rhythm deficits, nor histological changes in the suprachiasmatic nucleus of iTON mice. There was no difference in Fluoro-jade B staining (A, B), GFAP expression (C, D), or microglial soma area (E, F) at either 7 or 30 DPI in the SCN. (G) Injured mice were significantly less active than sham mice overall, but there was not a significant change in the circadian distribution of activity. Representative photomicrographs taken at 10x magnification. White dashed lines indicate region of interest/ area where measurements were taken. Scale bars indicate 100 µm.

### Injured mice have significantly increased axonal degeneration throughout the primary visual system

FJ-B staining for degeneration showed a pattern of punctate staining indicative of axonal degeneration throughout the majority of RGC projection targets [24, 45] (fig. 6). Areas examined were the optic tract (OT), superior colliculi (SC), ventral lateral geniculate nucleus (vLGN), dorsal lateral geniculate nucleus (dLGN), and suprachiasmatic nucleus (SCN). No FJ-B staining was present in the visual cortex of either sham or TON mice. Seven DPI TON mice present with increased degeneration (as calculated by mean fluorescence intensity) in the OT (TON n=8, sham n=10; t(16) = −9.4, p < 0.001), vLGN (TON n=7, sham n=7; t(11) = −3.1, p = 0.01), and dLGN (TON n =7, sham n=7; t(12) = −2.6, p = 0.02), and SC (TON n=7, sham n=8; t(13) = −6.0, p < 0001). There was no staining in the suprachiasmatic nucleus (SCN; TON n=8, sham n=7; p = 0.1; Fig. 5a). This degeneration is also present 30 DPI in OT (TON n=8, sham n=8; t(14) = −9.1, p < 0.001), vLGN (TON n=8, sham n=8; T = 42, p = 0.005), dLGN (TON n=8, sham n=8; t(14) = −4.4, p < 0.001), and SC (TON n=7, sham n=8; t(13) = −9.7, p < 0.001) but not SCN (TON n=8, sham n=6; t(12) = −0.76, p = 0.5; Fig. 5b). Interestingly vLGN and dLGN were also significantly different within TON mice (t(14) = 2.4, p = 0.03) with vLGN of injured mice having higher mean fluorescence intensity (*M*=0.99, *SD*=0.17) than dLGN (*M*=0.77, *SD*=0.17).

**Figure 6.**
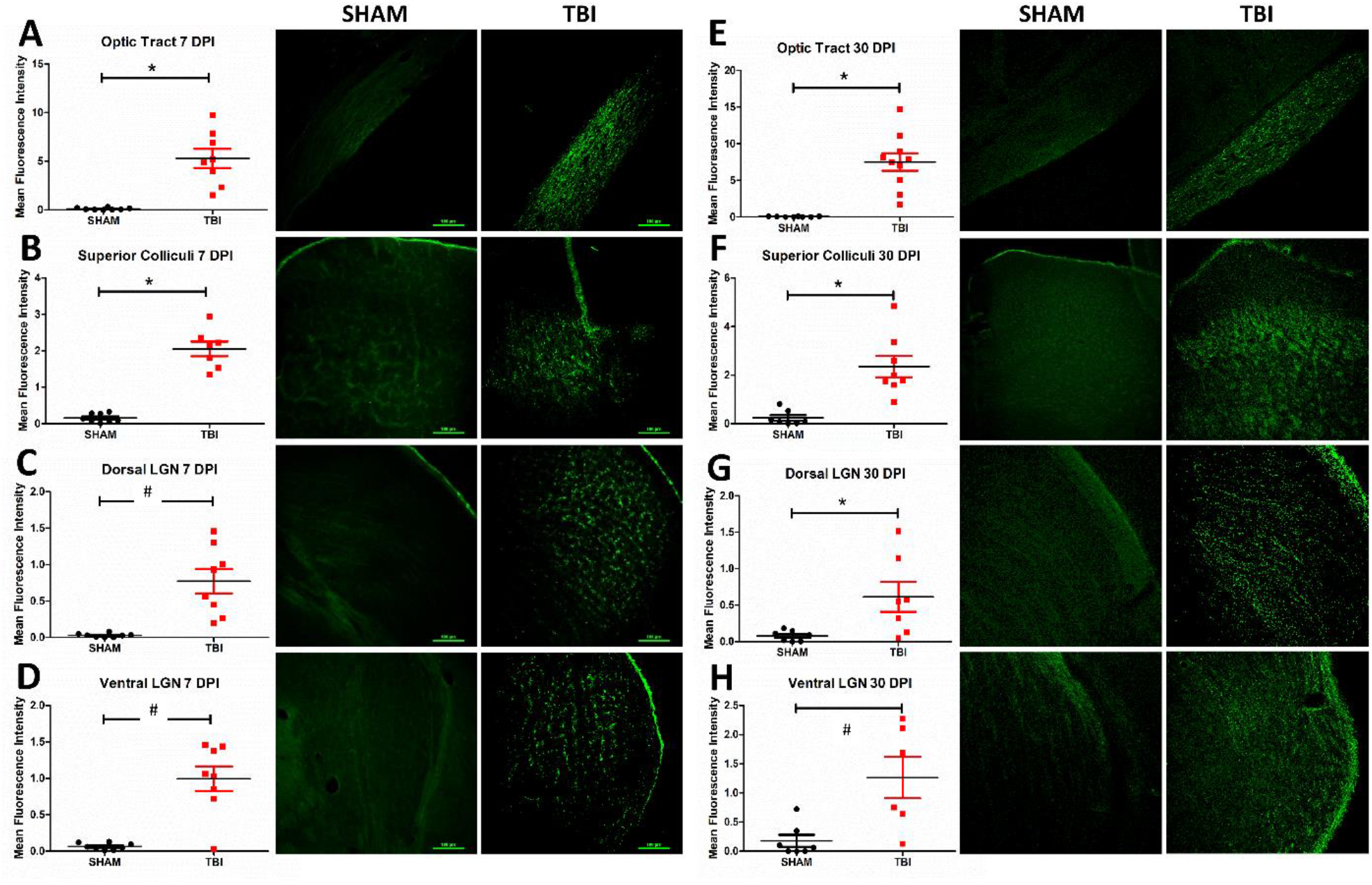
Neurodegeneration in the optic system 7 (A-D) and 30 (E-H) DPI. There was significantly elevated Fluoro-jade B staining in the optic tract (A,E), superior colliculus (B, F), dorsal lateral geniculate nucleus (LGN; C, G), and ventral LGN (D, H) at both 7 and 30 DPI in iTON mice compared shams. Staining appears punctate rather than somatic, consistent with primarily axonal staining. Representative photomicrographs taken at 10x magnification with scale bar indicating 100 μm. * p < 0.001, # p <0.05.

### There is microglial activation in some downstream optic subcortical thalamic targets

The same thalamic and brainstem targets of retinal ganglion cells were analyzed for microglial activation by Iba-1 fluorescent immunostaining 7 and 30 DPI (Fig. 7). Microglial soma area and perimeter were utilized to assess morphological changes in microglia. Seven DPI there was significantly increased soma area (TON n=8, sham n=8; t(13) = −4.5, p <0.001) and perimeter (t(13) = −4.2, p < 0.001) in OT, and area (TON n=9, sham n=6; t(13) = −2.12, p = 0.05) but not perimeter (p = 0.056) was significantly increased in dLGN. There was not a significant change in microglial morphology in the superior colliculi (area p = 0.056, perimeter p = 0.056), vLGN (area p=0.3, perimeter p= 0.7), or SCN (area p = 0.9, perimeter p = 1; Fig. 5e).

**Figure 7.**
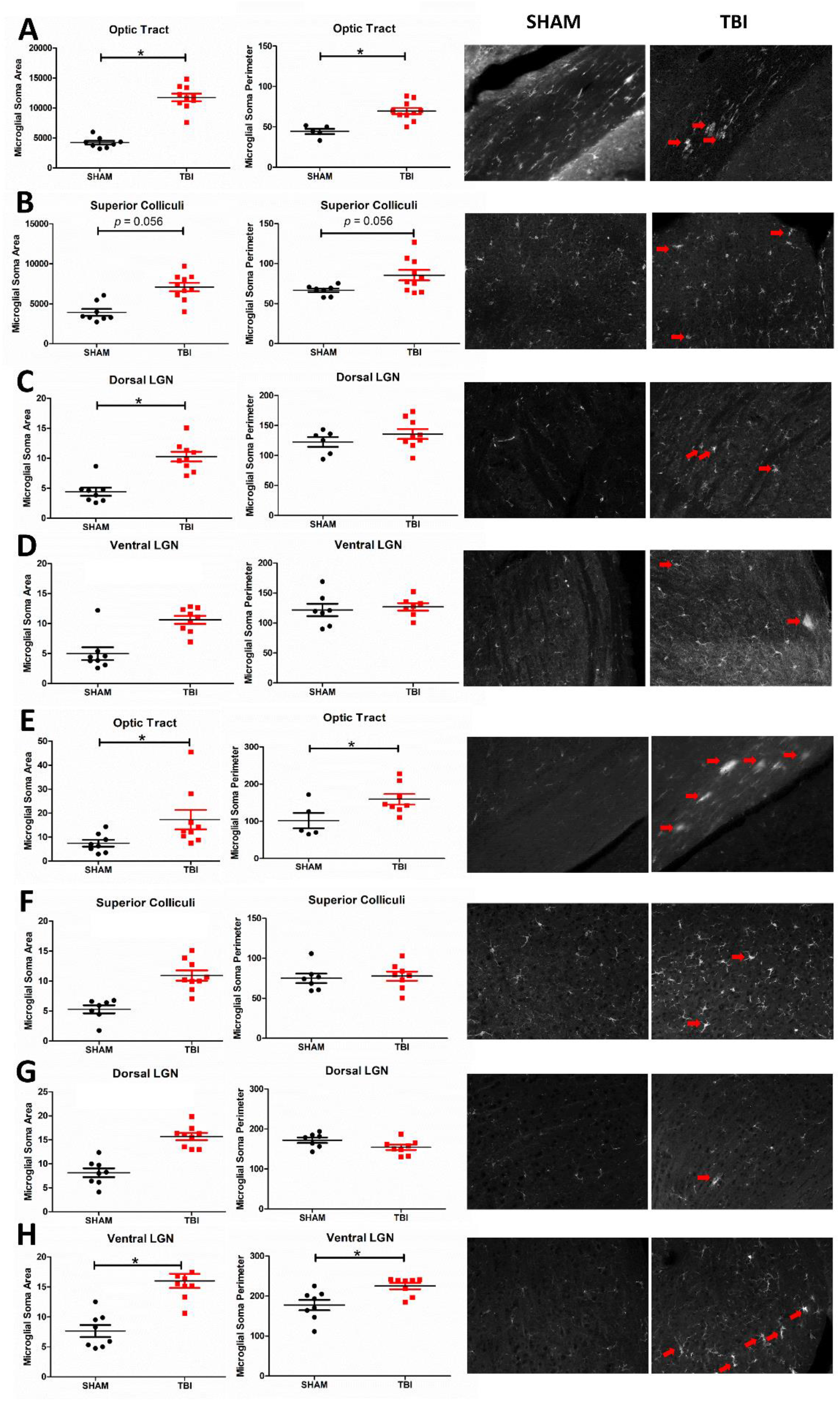
Morphologic evidence of microglial activation in the optic system is present 7 (A-D) and 30 (E-H) DPI. There were significant increases in microglial soma area (as measured using Iba-1 immunoreactivity) in the optic tract (A,E), Superior Colliculi (B, F), dorsal lateral geniculate nucleus (LGN; C, G), and ventral LGN (D, H) both 7 and 30 DPI, respectively, in iTON mice but not in shams. These morphologic changes are particularly predominant in the OT, but can also clearly be seen in other regions (red arrows). Representative photomicrographs taken at 10x magnification, scale bar 100 μm. * p < 0.001, # p <0.05

Thirty DPI, optic tract microglia are also significantly different from sham in area (TON n=8, sham n=5; t(19) = 4.1, p = 0.002) and perimeter (t(19) = 2.7, p = 0.04). vLGN area (TON n=8, sham n=8; t(22) = 4.0, p = 0.002) and perimeter (t(22) = 5.4, p = 0.01) are significantly higher in TON mice but not in the dLGN (area p=0.8, perimeter n=0.5). There was also not a significant change in microglial morphology in the superior colliculi (area p = 0.9 and perimeter p=0.8) or SCN (area: p = 0.9; perimeter: p = 0.8; Fig. 5f).

### There is astrogliosis in TON mice

Optic tract astrogliosis was also seen, with significantly elevated expression of GFAP 7 DPI (TON n = 8, sham n=8; t(16) = −9.8, p <0.001; Fig. 8). Moreover, there is reactive gliosis present in the vLGN (TON n=9, sham n=9; t(15) = −5.5, p < 0.001), dLGN (TON n=8, sham n=9; t(15) = −3.9, p = 0.001), and SC (TON n=8, sham n= 10; t(16)= −4.6, p < 0.001) of injured mice 7 DPI, but not in the SCN (TON n=5, sham n=6; p = 0.9; Fig.5c). These effects are also seen 30 DPI with significantly increased gliosis in the OT (TON n=8, sham n=8; t(14) = −2.6, p = 0.02), vLGN (TON n=8, sham n=9; t(15) = −5.3, p < 0.001), dLGN (TON n=8, sham n=9; t(15) = −6.4, p < 0.001), and SC (TON n=7, sham n=9; t(14) = −4.9, p < 0.001) of TBI mice but, again, not the SCN (p = 4.4; Fig. 5d).

**Figure 8.**
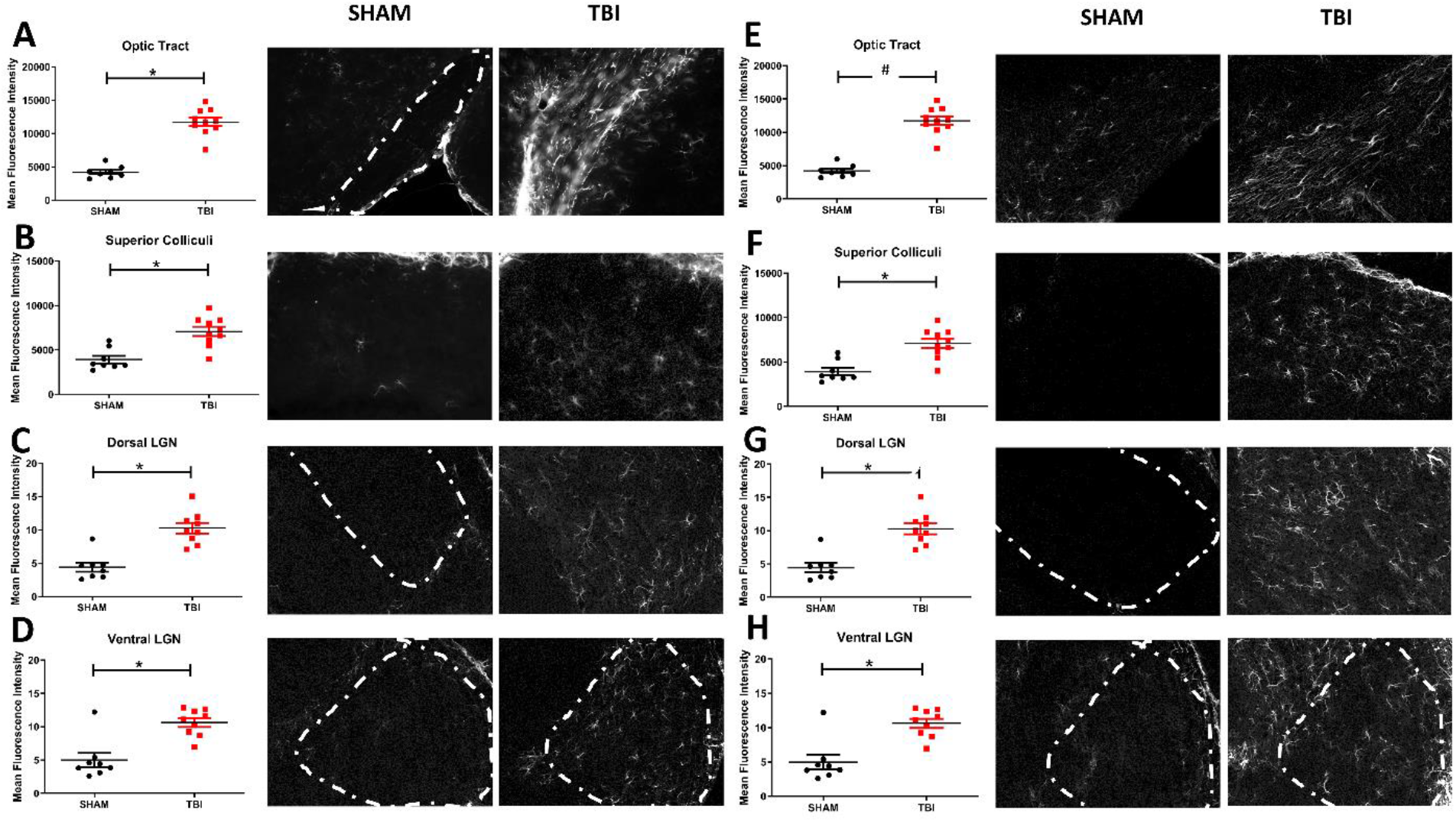
Astrogliosis of the optic system is significantly elevated in iTON mice 7 (A-D) and 30 (E-H) DPI as indicated by increased fluorescence intensity of GFAP. There was significant elevated GFAP intensity in the optic tract (A,E), superior colliculus (B, F), dorsal lateral geniculate nucleus (LGN; C, G), and ventral LGN (D, H) both 7 and 30 DPI in iTON mice as compared to controls. White dashed lines mark the areas analyzed in sham and iTON mice where it is more difficult to see visualize the location of the nuclei. Representative photomicrographs taken at 10x magnification, scale bar 100 μm. * p ≤ 0.001, # p <0.05

### ER stress markers are elevated after experimental head trauma

There are three major ER stress pathways in mammals. Two of these pathways function around the IRE-1α and PERK receptors with downstream effectors including PDI, CHOP, and eRO1L. We performed Western blotting on retinal protein extracted 7-and 30-days post injury. At seven DPI (Fig. 9) there were significantly elevated levels of IRE-1α (TON n=14, sham n=14; t(22) = −2.6, p = 0.02), PERK (TON n=14, sham n=11; T = 92, p = 0.006), ERO1-L (TON n=11, sham n=12; t(21) = −2.4, p = 0.02), and CHOP (TON n=17, sham n=9; t(24) = −5.7, p < 0.001), but not PDI (p = 0.4). At thirty DPI (Fig. 10), ER stress markers remain elevated in TON mice for IRE-1α (TON n=8, sham n=8; t(14) = 2.2, p = 0.04), ERO1-L (TON n=12, sham n=12; t(22) = −2.1, p = 0.04), and CHOP (TON n=8, sham n=8; t(14) = −2.5, p = 0.03). Interestingly, PERK activation is no longer significantly elevated (p=0.9), but PDI is now elevated (TON n=9, sham n=8; t(15) = 2.2, p = 0.04).

**Figure 9.**
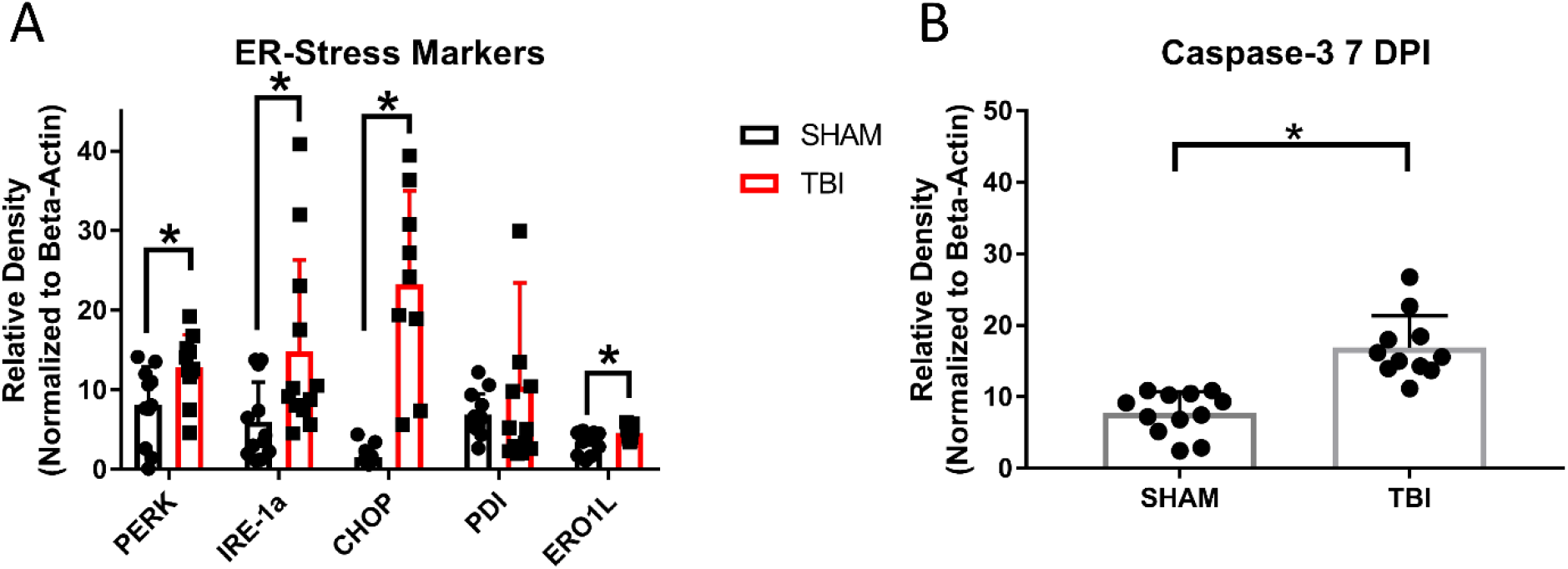
There is increased apoptosis and elevated markers of Endoplasmic Reticulum stress in the retinas of TBI/ iTON mice 7 DPI. Both ER stress receptors (IRE-1 and PERK) and downstream factors (CHOP) and well as ROS indicator (ERO1L) were elevated 7 DPI, while PDI was not (A). The apoptotic marker Caspase-3 was also significantly elevated in TBI mice. Protein measurements were normalized to beta-actin. *p < 0.05

**Figure 10.**
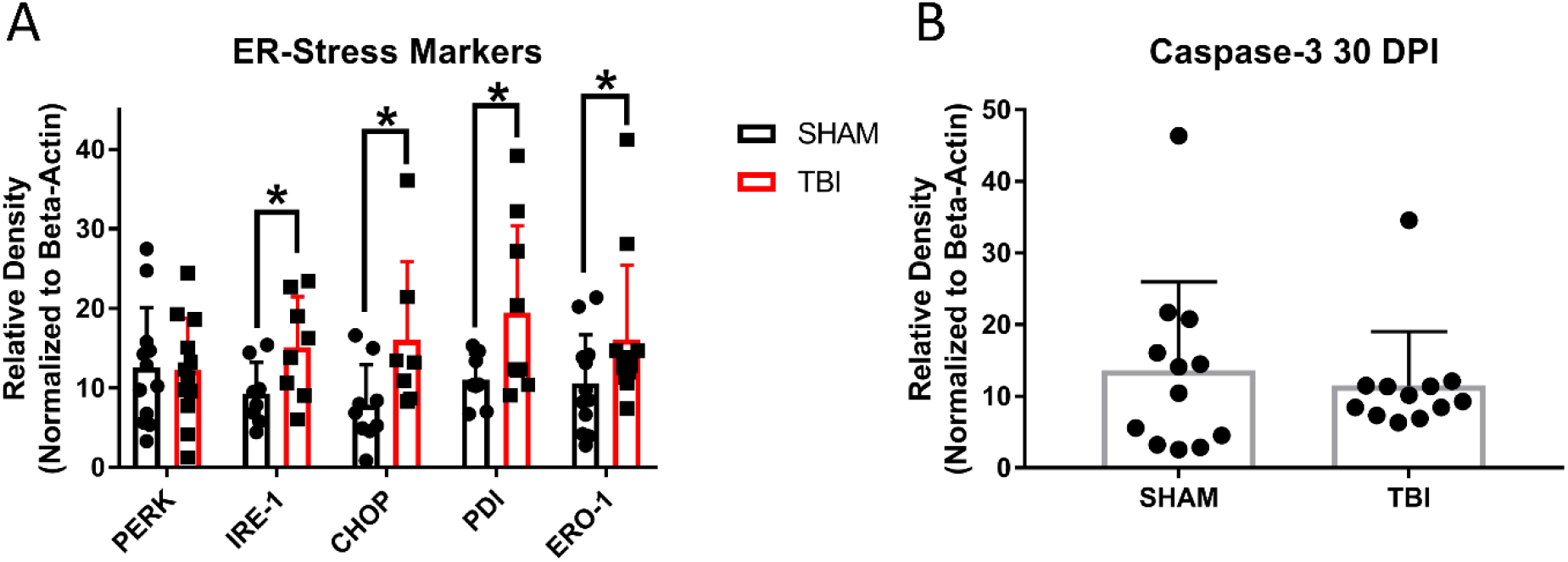
Elevated markers of Endoplasmic Reticulum stress are still present 30 DPI in TBI mice. (A) There is a change in the ER stress milieu of retinas 30 DPI such that expression of the IRE-1 receptor is elevated, but not PERK. CHOP expression is significantly higher in iTON mice, which may suggest that cells are still being directed toward apoptosis rather than protein refolding; however, caspase-3 expression is no longer significantly elevated (B). Interestingly, in contrast to the pattern seen earlier after injury, PDI is elevated at 30 DPI. *p < 0.05

## Discussion

Utilizing a mild weight-drop injury we have produced a novel model for analyzing TON within the common context of mild traumatic brain injury. This model recapitulates key features of TON without direct penetration of the optic nerve, is easily replicated, and shows evidence of injury from the molecular to the behavioral level for at least 30 days after injury. While we have previously shown similar results to those described here in adult mice,[24] the current study examines more features of TON and extends our findings to adolescent mice; thus, further validating this as a replicable model of TON across ages and multiple study groups. We have shown that indirect TON results in injury not only to the optic nerve, with associated retinal ganglion cell death, but also shows effects in more downstream subcortical and brainstem targets of RGC axons including the dorsal and ventral lateral geniculate nuclei and superior colliculi, but not the suprachiasmatic nucleus or visual cortex. Lack of differences in circadian activity between sham and injured mice over the first 11 DPI is consistent with absence of histologic damage in the SCN through 30 DPI. We have additionally provided evidence that this injury affects the optokinetic response and that degeneration, astrogliosis, and microglial activation are accompanied by elevated endoplasmic reticulum stress markers up to 30 days post injury.

We first attempted to extend our previous findings by adding a behavioral measure of impaired visual function. We, therefore, exploited a common behavioral assay used in models of optic degeneration, retinitis pigmentosa, and other optic neuropathies for the optokinetic response, which is correlated with visual acuity.[39] TON mice both 7 and 30 DPI experience a significantly blunted OKR compared to their uninjured counterparts. It is so blunted that injured mice only elicit the OKR 1-3 times at their “optimal visual spatial frequency,” which was determined to be between .13 and .26 cpd in C57Bl/6 mice by our own pilot data and that of Abdeljalil, et al.,[42] throughout the four-minute recording period compared to roughly 12-14 times in their sham counterparts (consistent with previous reports that C57Bl6 mice have normal vision).[42] The OKR is a critical visual function as it allows one to perceive and follow objects as they cross the field of vision as well as the proper function of smooth eye movements and other reflex eye movements.[46] OKR deficits could also indicate a decline in visual acuity as one needs to be able to see an object to follow it.[39] Tao et al. utilized optical coherence tomography and flash electroretinograms to argue that there is both a thinning of the retinal layers and reduced RGC functioning respectively in an ultrasonic, indirect TON model.[25] Our results now add that there is decreased OKR and smooth pursuit functioning and possibly reduced visual acuity, since we see no response in injured mice at the highest spatial frequency of 0.46, which sham animals were still responding to.

Additionally, while we cannot directly correlate the 7-and 30-day groups because the assays were run in separate cohorts, there appears to be subjective recovery of the optomotor response at 30 DPI as compared to at 7 DPI. This is consistent with the finding of visual system plasticity leading to visual recovery in a fluid percussion TBI model that also features optic nerve injury.[47] Of additional note, blinded scorers of the OKR assay made note of several behaviors in injured mice that were not noted in control mice. While these behaviors were not the focus of this paper, they may provide further avenues of research in the future. For example, compared to shams, injured mice tended to spin in circles opposite to the direction of the spinning drum and froze for longer periods. This could be related to motion and light sensitivity which are commonly reported in patients with TBI,[48, 49] suggesting that future research examining these behaviors may be useful for research into other types of visual deficits after head trauma. Interestingly, this is one of only a handful of times the OKR has been used as an outcome measure in TBI models.[50]

In addition to behavioral vision deficits, we also show a significant reduction in the number of retinal ganglion cells of injured mice both 7 and 30 DPI. Seven days post injury there were significantly reduced RGCs across all three retinal regions analyzed, but at 30 DPI this was only true in outer and middle regions of injured mice. Our analysis of the Caspase 3 apoptotic marker suggests the presence of apoptotic cells early after injury but not at 30 DPI. Thus, RGC death is an early consequence of this indirect injury that results in RGC apoptosis. The continued presence of cells expressing apoptotic markers at 7 DPI may suggest a window of several days after injury for possible therapeutic interventions.

Our data are consistent with RGC loss due to degeneration, which is initiated by injury to the intracanalicular portion of the ON.[24] Positive FJ-B staining in the ON is present 7 and 30 DPI. Of note, the pattern of histologic staining for degeneration seen in these experiments (Fig. 6) is punctate, and morphologically consistent with axons. This suggests that there are degenerating axons in the regions noted. Supporting evidence for similar axonal injury in the optic nerve has also been found in blast models [23-33, 34] and midline fluid percussion injury as well as targeted DIA to the optic nerve [47]. We did not see evidence of trans-synaptic neuron loss as there was no evidence of cell soma degeneration in brain areas with trans-synaptic optic nerve projections (e.g. visual cortex). We have also shown that markers of neuroinflammation and astrogliosis are elevated 7 and 30 DPI in the optic nerve/tract, suggesting that the pathologic processes have not yet resolved at these times. These results suggest that degenerative processes may persist for long periods after injury. These long-term effects have been reported previously in both indirect and direct models of optic nerve injury,[25, 51] although long-term functional consequences of this are not yet fully elucidated. It has also been shown that gliosis surrounding axonal damage likely progresses along the axons to its far-reaching projections.[52] Thus, we examined multiple projection targets of the optic nerve for evidence of degeneration.

Second, there is increased degeneration and gliosis, but not microglial activation, in the superior colliculi 7 and 30 DPI. These data indicate that the axonal degeneration seen in the anterior optic tract extends to the midbrain colliculi. Further, because the SC plays a role in the integration of visual attention, saccadic eye movement, and localization of attention shifts to new stimuli, the deficits in OKR performance may be explained in part by these histological findings. Future studies should more closely examine other areas involved in the OKR like those of the accessory optic system as well as attempt to test these distinct visual functions. The fact that there is no microglia reactivity 7 nor 30 DPI suggests that the pathophysiology of injury here may be driven more by reactive gliosis rather than neuroinflammation. Recent studies probing the timing of astrocytic versus microglial activation following TBI have suggested that microglia are present closer to the site of injury initially while downstream locations favor astrocyte activation with microglial inflammation occurring even years after the initial insult.[53] Thus, it is possible that the difference in microglial morphologic changes between the LGN and SC could be due to the fact that the SC is physically further from the site of injury.

We also visualized neurodegeneration, gliosis, and activated microglia in thalamic projections of the ON in the LGN. The LGN is anatomically divided into ventral and dorsal sub-nuclei. This division is important because the vLGN is more directly connected to the superior colliculi and has also been implicated in the optokinetic response and circadian rhythm, while the dLGN is known to project primarily to the visual cortex in both rats and cats.[54, 55] At seven DPI, injured mice have significant axonal degeneration, indicated by positive, punctate FJ-B staining, in both vLGN and dLGN, though they are not significantly different from each other. Thirty DPI both vLGN and dLGN also have significantly greater degeneration than found in sham animals, and the vLGN has significantly higher FJ-B intensity than the dLGN in TON mice. While we are uncertain as to why the two regions differ at 30 days, this finding is persistent across all measures taken (elaborated on below), such that the projections to the dLGN and vLGN appear to be responding to this injury differently, and this could provide an interesting avenue for future research. Additionally, lack of degeneration in the visual cortex despite positive staining in the dLGN may indicate that this injury is not (yet) trans-synaptic at the time points analyzed.

Many studies of TBI, axonal injury, and/or direct TON have shown increased microglial activation,[53-56-59] but this has focused on the injury site or optic nerve itself. Few studies have addressed neuroinflammatory changes in downstream projection targets. We have shown that seven DPI there is significantly increased soma area of microglia in the dLGN but not the vLGN. Thirty DPI, however, there is significantly changed microglial morphology in the vLGN but not the dLGN. The reason for this reversal is not clear from the present studies. One possibility may be differences in susceptibility of RGC subtypes to this injury, as this has been shown in other optic nerve models.[60, 61] Moreover, there is increased astrogliosis in both vLGN and dLGN both seven and thirty DPI. This may suggest that astroglia are playing a larger role in the detrimental effects of the injury subacutely than the neuroinflammation implied by morphologically activated microglia.

Prolonged elevation of astrocyte activity as well as persistent microglial activation can result in protective or pernicious outcomes, or both. It has recently been disputed whether gliosis serves more immediate protective functions through the formation of a gliotic scar at the site of injury; or through the release of growth factors (TGFβ) and cytokines (IL-6), that support repair of injured tissue and reduce neuroinflammation; or even by engulfing the injured tissue and inhibiting contact-induced apoptosis[58-62, 63] or if the consequences of activation are more deleterious. For example, the gliotic scar prevents axonal regrowth and repair,[58] but removal of proliferating glia (i.e., glia that form the gliotic scar) results in increased neurodegeneration to the impact site, so astrocytes’ roles are not yet fully understood.[64] Still other research suggests that it is the milieu of microglia and astrocytes together that is critical for proper improvement after injury. Microglia are signaled in response to gliosis but can also respond to injury in more acute periods, preceding development of the gliotic scar.[53] Fitch and Silver further argue that proximity to the injury may determine the dichotomous roles of astrocytes such that those inhibiting axon regrowth may be closer to the injury and those releasing beneficial growth factors may be more distant.[52] Identifying activated astrocytes by GFAP content, as was done in this study, would not differentiate these activities, so future studies could more specifically differentiate between these two possibilities.

Though the vLGN is implicated in circadian rhythm regulation[65] there were no indications of circadian dysfunction in this injury model. There was also no evidence of degeneration, inflammation, or gliosis in the SCN, nor were there any shifts in activity patterns measured over 11 DPI despite animal TBI studies showing shifts in both motor activity and orexin levels 3 days post injury [66], as well as acute loss of cortisol-mediated circadian rhythm in adults 18-65 [67], and increased melatonin-mediated shifts in children after severe TBI [68]. It is also possible, though, that circadian shifts may take longer to develop as has been true in human populations ranging from 4 months to 2 years after injury [68]. This may indicate that a subset of RGCs, known as intrinsically photosensitive RGCs, that project directly to the SCN and help mediate light entrainment of circadian rhythms,[69] may be less susceptible to injury in this model. Indeed, this is the case in optic nerve crush and transection models.[60] It is also possible that this difference in apparent injury susceptibility between RGC populations is due to the part of the cross-sectional area of the optic nerve in which these fibers run, since in at least some species there is consistent, likely retinotopic, organization of optic nerve fibers within the nerve.[70] In this case, some fibers in the ON could possibly be protected by their relative position in the nerve (for example, by being in the center of the nerve and thus partially shielded from impact). However, in our studies degenerating axons appear to be distributed throughout the optic tract diameter (Fig. 6a/e) rather than having a discernible geometric pattern within the tract.

There is a significant body of literature on the effects of ER stress on many models of optic neuropathies.[14] In spite of this, relatively little is known about the role of ER stress in the pathology of TON associated with head trauma. We analyzed retinas of sham and injured mice to determine to what degree major ER stress pathways were activated by the experimental TON. Indeed, two of the three major pathways (IRE-1α and PERK) were upregulated seven DPI. Additionally, the downstream apoptotic effector of the ER stress pathway – CHOP – was upregulated in TON mice 7 DPI. This suggests that after injury, cellular stress is high, and the unfolded protein response appears to lead to apoptosis rather than refolding even in the acute phase after TBI.[71] This interpretation is supported by our finding of apoptotic Caspase 3 upregulation in retinas of TON animals.

Interestingly, PDI is not changed 7 DPI. PDI has two functions in the ER stress pathway – as a redox-sensing activator of IRE-1α and PERK receptors and as a downstream nuclear chaperone of the UPR PERK pathway that promotes refolding of misfolded proteins.[71-73] This lack of PDI upregulation relatively early after injury may indicate that it is not acting as an ER stress agonist in this injury model and/or that the level of stress is too high for the cell to push for repair over apoptosis. It could also be that an acute period of activation occurs prior to our earliest measurements at 7 DPI. This is supported by the finding of apoptotic CHOP and Caspase3 elevation. The elevation of ERO-1L further supports the conclusion of elevated ER stress, and also implicates oxidative stress, since this protein is active in both oxidative stress and ER stress pathways.[74] Veritably, recent literature suggests oxidative and ER stress pathways are interconnected, although mechanisms are not fully clear, at least within TBI studies.[14-75-77]

Thirty DPI IRE-1α, CHOP, and ERO-1L are elevated in TON mice. In contrast to the 7 DPI cohort, PERK is not significantly elevated, but PDI is. The delayed increase in PDI favors the function of PDI in this injury model as a chaperone for protein repair and may indicate potential for therapeutic strategies within a month post injury where RGCs are no longer dying but recovering from the injury. This potential shift to a repair response rather than apoptosis is supported by the lack of significant difference in Caspase 3 expression between sham and TON mice 30 DPI, which suggests that the apoptotic response has subsided even though CHOP expression is still high. The concurrent and persistently elevated ERO-1 and IRE-1 may also indicate that redox stress is also long-lasting and PDI may be acting to keep IRE-1α activated[72] while PERK is not (Fig. 10a). Future studies would need to find a way to separate these stress responses and determine whether these proteins are working through united or separate mechanisms.

## Conclusions

In summary, we show that this closed head trauma model in adolescent mice leads to reproducible optic tract injury, and this can be used as a model of indirect traumatic optic neuropathy. We describe significant changes to the optokinetic response that are present early and late after injury. There is clear RGC loss 7 and 30 DPI and active apoptosis 7 DPI. There is axonal degeneration, neuroinflammation, and gliosis in ON, SC, and LGN. The elevation of multiple early and late ER stress markers suggests that ER stress is a possible pathologic mechanism for the ongoing degeneration. There is also evidence of both ER stress and oxidative stress through persistent ERO-1L elevation. This model shows promise for ongoing study into the pathologic mechanisms and potential treatment strategies for indirect TON associated with head trauma.

## Acknowledgements

We thank Jordyn Torrens and Emily Shalosky for assistance in performing the studies described. This work was supported by the National Institutes of Health [NIH grant HD001097 (NKE)], and by a Procter Award and the Division of Pediatric Rehabilitation Medicine at Cincinnati Children’s Hospital Medical Center (NKE). Funding entities played no role in the design, execution, analysis, or interpretation of results.

## Author Contributions

All authors had full access to all the data in the study and take responsibility for the integrity of the data and the accuracy of the data analysis. *Conceptualization:* S.M.C., F.G.C., and N.K.E.; *Methodology: S*.*M*.*C, F*.*G*.*C*., *D. D*., *and N*.*K*.*E*.; *Investigation:* S.M.C., F.G.C, and A. M. B., and D. D..; *Formal Analysis:* S.M.C, F.G.C., A. M. B. and N.K.E.; *Resources:* N.K.E.; *Writing – Original Draft:* S. M. C., F.G.C., A.M.B., D.D, and N.K.E; *Writing – Review & Editing:* F.G.C., S.M.C, A.M.B, D.D., and N.K.E; *Visualization*: F.G.C., S.M.C., D.D., and N.K.E.; *Supervision*: N.K.E.; *Project Administration:* F.G.C, S.M.C., and N.K.E.; *Funding Acquisition:* N.K.E.

## Data Accessibility

The data that support the findings of this study are available from the corresponding author upon reasonable request.

